# Gastrointestinal carriage is a major reservoir of *K. pneumoniae* infection in intensive care patients

**DOI:** 10.1101/096446

**Authors:** Claire L Gorrie, Mirjana Mirceta, Ryan R Wick, David J Edwards, Richard A Strugnell, Nigel Pratt, Jill Garlick, Kerrie Watson, David Pilcher, Steven McGloughlin, Denis W Spelman, Adam W J Jenney, Kathryn E Holt

## Abstract

**Background:** *Klebsiella pneumoniae* is an opportunistic pathogen and a leading cause of hospital-associated (HA) infections. Patients in intensive care units (ICUs) are particularly at risk, and outbreaks are frequently reported in ICUs. *K. pneumoniae* is also part of the healthy human microbiome, providing a potential reservoir for HA infection. However, the frequency of *K. pneumoniae* gut colonization and its contribution to HA infections are not well characterized.

**Methods:** We conducted one-year prospective cohort study of ICU patients. Participants (n=498) were screened for rectal and throat carriage of *K. pneumoniae* shortly after admission, and clinical information was extracted from hospital records.*K. pneumoniae* isolated from screening swabs and clinical diagnostic samples were characterized using whole genome sequencing. Genomic and epidemiological data were combined to identify likely transmission events.

**Results and Conclusions:** *K. pneumoniae* carriage frequencies were estimated at 6% (95% CI, 3%-8%) amongst ICU patients admitted direct from the community, and 19% (95% CI, 14% – 51%) amongst those who had recent contact with healthcare. Gut colonisation on admission was significantly associated with subsequent *K. pneumoniae* infection (infection risk 16% vs 3%, OR=6.9, p<0.001), and genome data indicated a match between carriage and infection isolates in most patients. Five likely transmission chains were identified, resulting in six infections (12% of *K. pneumoniae* infections in ICU). In contrast, 49% of *K. pneumoniae* infections were caused by a strain that was unique to the patient, and 48% of patients with *K. pneumoniae* infections who participated in screening were positive for prior colonisation. These data confirm *K. pneumoniae* colonisation is a significant risk factor for subsequent infection in ICU, and indicate that half of all *K. pneumoniae* infections result from patients’ own microbiota. Screening for colonisation on admission could limit risk of infection in the colonised patient and others.

## Introduction

*Klebsiella pneumoniae* is a significant cause of morbidity and mortality worldwide and causes a range of opportunistic infections in susceptible hosts including the young, elderly, immunocompromised, hospitalized and particularly those in intensive care units (ICUs)^1–3^. Infections with *K. pneumoniae* are increasingly refractory to antibiotic treatment due to rising rates of antibiotic resistance^4–6^, and *K. pneumoniae* is amongst the ESKAPE pathogens responsible for the majority of hospital-associated (HA) infections^7^. The US Centre for Disease Control and Prevention (CDC) have identified carbapenemase-producing (CP) *Enterobacteriaceae* and extended spectrum beta-lactamase (ESBL) producing *Enterobacteriaceae* as significant threats to global health, and found *Klebsiella* species comprised 11% and 23% of healthcare-associated CP and ESBL infections in the US, respectively^4^. The emergence of multidrug resistant (MDR) *K. pneumoniae* has resulted in a dramatic increase in research into reservoirs and risk factors for HA *K. pneumoniae* infections, largely focused on ESBL or CP bacteria isolated from infections and intra-hospital outbreaks. These studies have demonstrated transmission of ESBL or CP *K. pneumoniae* between patients, and show that gastrointestinal (GI) tract colonisation with ESBL or CP *K. pneumoniae* can be a risk factor for infection^6,8^. Yet whilst the majority of *K. pneumoniae* HA infections are not ESBL or CP^3,9^, there is little data on the frequency and clinical relevance of colonisation with *K. pneumoniae* more generally.

*K. pneumoniae* is known to asymptomatically colonise the skin, mouth, respiratory and GI tracts of humans. Analysis of 16S rRNA gene data from healthy volunteers in the human microbiome project detected *K. pneumoniae* in approximately 10% of samples from the mouth, nares and skin, and 3.8% of stool samples^10^. Two recent studies investigated carriage of *K. pneumoniae* in the respiratory tracts of healthy individuals in communities in Asia, using bacteriological culture. A 2010 study in Indonesia detected nasopharyngeal carriage in 15% of adults and 7% of children^11^, while a 2014 study in Vietnam detected nasopharyngeal carriage in 2.7% of adults and throat carriage in 14%^12^. In addition, it has recently been shown that patients suffering from CP *K. pneumoniae* infections (typically clonal group 258), or from pyogenic liver abscess caused by hypervirulent *K. pneumoniae* (typically clonal group 23), frequently carry their infecting strain in their GI tract for between 30 days (≤74%) and six months (<30%) following discharge from hospital^13^.

If the gut microbiome is a common reservoir for HA *K. pneumoniae* infection, then the risk of such infections could potentially be mitigated through screening and GI decontamination^14^, as is often done for vancomycin resistant *Enterococcus* (VRE). The question of whether *K. pneumoniae* colonisation poses a risk for subsequent infection has been investigated in just two studies to date. A 1971 study found 18.5% of patients admitted to various wards in the Denver Veterans Administration Hospital were positive by culture for rectal carriage of *K. pneumoniae*, and that carriage was significantly associated with risk of subsequent HA infection (45% vs 11%)^15^. A 2016 study at the University of Michigan Health System tertiary care hospital reported similar colonisation rates (23%) and increased risk of infection following colonisation (5.2% in colonised vs 1.3% in non-colonised)^16^.

In the present study, we assessed the prevalence of *K. pneumoniae* colonisation in an at-risk cohort in an ICU within a modern, well-equipped, and well-managed tertiary teaching hospital in Australia. Additionally, we investigated whether colonisation on admission enhances risk of subsequent *K. pneumoniae* infection among ICU patients and the relative contribution of patients’ own gut microbiota and intra-hospital transmission to the burden of *K. pneumoniae* carriage and infection in the ICU.

## Methods

### Ethics

Ethical approvals for these studies were granted by the Alfred Hospital Ethics Committee (Project numbers #550/12 and #526/13).

### Setting and recruitment

The *Klebsiella* Acquisition Surveillance Project at Alfred Health (KASPAH) was conducted from April 1, 2013 to March 31, 2014. The study included monitoring for all clinical isolates identified as *K. pneumoniae* infections by the hospital diagnostic laboratory (details in **Supplementary Methods**) and recruitment of adult patients (≥18 years) in the Alfred Hospital ICU for *K. pneumoniae* carriage screening via rectal and throat swabs. Verbal consent was needed from patients (or an adult responsible for the patient) for the first nine months (consent-based collection). For the last three months (universal collection) a MDR surveillance study (#526/13) was concurrently conducted in the ICU by our group and did not require verbal consent. Additional details are provided in **Supplementary Methods**. As no differences were observed in the distributions of age, sex or *K. pneumoniae* carriage between the patients enrolled during consent-based and universal collection protocols (Supplementary Figure 1), results from these two groups were combined for all analyses.

### Sample and data collection

Baseline screening swabs and accompanying patient data was collected for all study participants as soon as possible after recruitment. Follow-up swabs and data collection were repeated each 5-7 days after baseline for the duration of ICU stay and up to 4 days following transfer to another ward. Details of swabbing procedures, bacterial culture and antimicrobial susceptibility testing are given in **Supplementary Methods**. Briefly, samples were grown on selective media and any with the appearance of *Klebsiella* species were investigated using matrix-assisted laser desorption ionization-time of flight (MALDI-TOF) and subjected to antimicrobial profiling using Vitek2. Information on age, gender, dates of hospital and ICU admission/s, surgery in the last 30 days, and antibiotic treatment in the last 7 days were extracted from hospital records. Dates of discharge and/or death were extracted from hospital records at the conclusion of the study. All clinical isolates recovered from ICU patients and identified as *K. pneumoniae* infections by the hospital diagnostic laboratory as part of routine care were included in the study.

### Community associated vs healthcare associated carriage

Individuals in the community associated (CA) screening group include patients who were both (i) admitted to the Alfred Hospital ICU either directly (day 0) or via another ward on day 0, 1 or 2 of the original Hospital admission; and (ii) first swabbed on day 0, 1 or 2 of that admission. Patients first swabbed on day 3 or later of their Hospital admission are included in the HA/Day 3+ screening group. Individuals referred to the Alfred Hospital ICU by the trauma ward of another hospital were assumed to be emergency admissions from the community, and were assigned to the CA/Day 0-2 or HA/Day 3+ screening groups according to the day of first swab relative to their Alfred Hospital admission. All other patients transferred from another hospital were included in the HA/D3+ screening group. As such, the CA/Day 0-2 screening groups represent individuals admitted directly to the hospital from the community, whereas the HA/D3+ group includes individuals with recent hospital exposure.

### DNA extraction and sequencing

DNA was extracted from overnight cultures using a phenol:chloroform protocol and phase lock gel tubes and sequenced via Illumina HiSeq to generate 125 bp paired-end reads (see **Supplementary Methods**). Following quality control checks of the sequence data, 148 isolates from 106 patients were subjected to comparative genomic analysis. Data and accession numbers on all isolates included are available in Supplementary Table 1. A maximum likelihood phylogenetic tree was inferred from an alignment of all SNPs identified within core *K. pneumoniae* genes using FastTree v2.1.8 ^17,18^. Lineages were defined based on this tree using RAMI^19^ and multi-locus sequence types were assigned using SRST2^20^. Isolates falling within the same lineage were further investigated to identify pairwise SNPs via assembly and read mapping. Full details of genomic analyses are given in **Supplementary Methods.**

### Statistical analysis

All statistical analyses were conducted using R (v3.3.1), including Fisher’s exact test for association and least squares linear regression for modelling the relationship between pairwise genetic distance and time between isolation (details in **Supplementary Methods**).

## Results

### *K. pneumoniae* carriage

A total of 498 ICU patients were recruited and screened for *K. pneumoniae* carriage. This represents one fifth of all ICU admissions in the study period, and 64% of adults spending more than 72 hours in ICU. Fifty-four patients (10.8%) tested positive at baseline screening (50 GI carriage only, 2 throat carriage only, 2 both). Carriage was detected at the same frequency in males and females (11.0% vs 10.4%; p = 0.9) and there was a trend towards higher carriage rates among older individuals (Supplementary Table 2).

We estimated the rate of CA *K. pneumoniae* GI carriage, among patients recruited and swabbed in the ICU within two days of their first recorded admission to the Hospital (CA/D0-2 group, see **Methods**), to be 5.9% (95% confidence interval (CI) [3% - 8%], Table 1, Supplementary Table 3). The HA GI carriage rate, amongst patients who were first swabbed in the ICU on or after the third day of admission to the Alfred hospital or following referral from another hospital (HA/D3+ group, see **Methods**), was significantly higher at 19% (95% CI [13.6% – 25.7%], OR=3.75, p=0.00001 using Fisher’s exact test, Table 1).

**Table 1.** *K. pneumoniae* GI carriage detected at baseline screening of Alfred Hospital patients. Patient groups: ICU CA/Day 0-2, rectal screening swab obtained on day 0, 1 or 2 of admission to Alfred Hospital and not referred from another hospital (except from trauma unit); ICU HA/Day 3+, rectal screening swab obtained on day 3 or later of admission to Alfred Hospital or referred from another hospital.

| Patient group | Total screened | *K. pneumoniae* screening swab result |  |
| --- | --- | --- | --- |
|  |  | Culture positive (%) | MDR (%) |
| ICU |  |  |  |
| CA / Day 0-2 | 324 | 19 (5.9%) | 0 (0%) |
| HA / Day 3+ | 174 | 33 (19.0%) | 6 (17.6%) |
| Total | 498 | 52 (10.4%) | 6 (11.3%) |

One third of the participants in the ICU screening study (n=170) contributed one or more follow-up screening swabs (Table 2). The overall GI carriage rate at follow-up was 15.3% (n=26/170), similar to the HA GI carriage rate of 19% (95% CI [0.71 – 2.38], OR=1.3, p=0.39). Participants testing positive on follow-up rectal swabs included 19 who tested negative for *K. pneumoniae* on their baseline rectal swab, yielding a conversion rate of 12%.

**Table 2.**
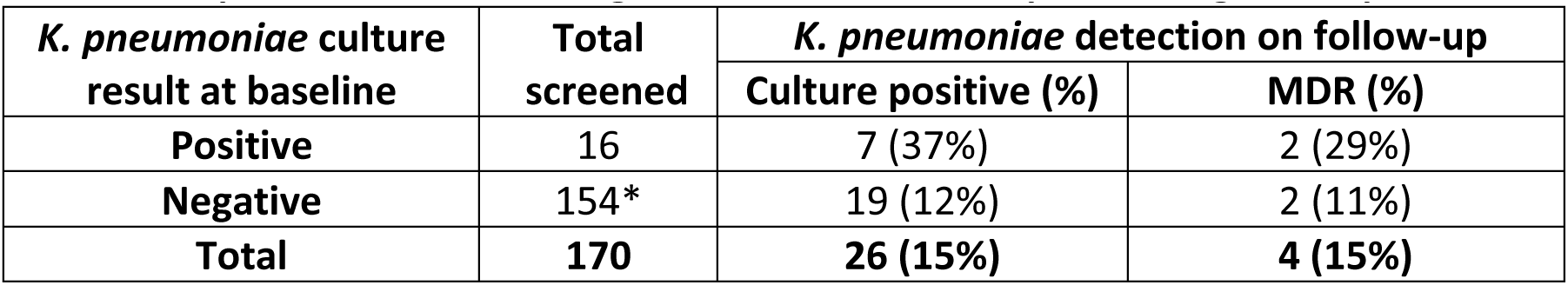
*K. pneumoniae* GI carriage detected in follow-up screening of ICU patients.

| *K. pneumoniae* culture result at baseline | Total screened | *K. pneumoniae* detection on follow-up |  |
| --- | --- | --- | --- |
|  |  | Culture positive (%) | MDR (%) |
| Positive | 16 | 7 (37%) | 2 (29%) |
| Negative | 154* | 19 (12%) | 2 (11%) |
| Total | 170 | 26 (15%) | 4 (15%) |

\*2 patients here were typed as *K. pneumoniae* follow-up swab-positive but WGS later showed these isolates as *E. coli*, and are not counted in the 19 follow-up positive.

None of the 19 CA baseline carriage isolates were MDR (Table 1). GI carriage of MDR strains was detected at similar rates among HA baseline isolates (18%, including 4 ESBL and 2 CP isolates, all in patients who had received antibiotics in the last 7 days) and follow-up screening isolates (16% of patients, including 4 with ESBL and 1 with CP isolates). A total of seven patients contributed both baseline and follow-up GI carriage isolates (Table 2). For 4/7 patients the resistance profiles for the follow-up isolate remained the same as the baseline isolate, for 2/7 patients the follow-up isolate was more resistant (one had increased resistance to amoxicillin/clavulanic acid, ticarcillin/clavulanic acid, piperacillin/ tazobactam, ceftazidine and decreased resistance to nitrofurantoin; the other had increased resistance to cefoxitin and nitrofurantoin), and for 1/7 the follow-up isolate was less resistant (decreased resistance to nitrofurantoin).

### *Klebsiella* GI colonisation is a source of infection in ICU

A total of 49 patients (1.8% of all adult ICU admissions) who spent time in the ICU during their hospital stay, were identified as having *K. pneumoniae* infections (11 ESBL, of which 3 were also CP). Most of these patients (n=39) were in the ICU when the *K. pneumoniae*-positive diagnostic specimen was taken, and ten were in another ward shortly after transfer from the ICU. Pneumonia was the most frequent form of *K. pneumoniae* infection in ICU patients (60%), followed by wound infections (15%), non-disseminated UTI (10%) and bacteraemia with sepsis (8%). MDR was most common among wound and blood isolates (>50%) (Table 3).

**Table 3.** Patients with infection(s) and time in the ICU. Type of infection and source of infection outlined (position/presence in transmission chain, prior colonisation, unknown). Note 3 patients had UTI and bacteraemia with sepsis; they are represented here in the bacteraemia with sepsis column.

|  | Pneumonia | UTI (non-invasive) | Wound | Other | Bacteraemia with sepsis | Total |
| --- | --- | --- | --- | --- | --- | --- |
| Recipient in transmission chain | 2 | 1 | 2 | 0 | 1 | 6 |
| Donor in transmission chain | 3 | 0 | 1 | 0 | 0 | 4 |
| Prior GI colonisation | 6 | 1 | 1 | 0 | 0 | 8 |
| Prior throat colonisation | 1 | 0 | 0 | 0 | 0 | 1 |
| Unknown source (unique lineage) | 17 (12) | 3 (3) | 3 (1) | 3 (2) | 3 (1) | 29 |
| Total | 29 (5 MDR) | 5 | 7 (4 MDR) | 3 | 4 (2 MDR) | 48 |

In order to assess whether *K. pneumoniae* GI carriage on admission to ICU was a risk factor for subsequent *K. pneumoniae* infection during hospital stay, we examined the subset of 491 individuals whose baseline screening swab was obtained at least two days prior to collection of any clinical specimen from which *K. pneumoniae* was isolated (Figure 1). The rate of *K. pneumoniae* infection was significantly higher amongst patients who were culture-positive for GI carriage at baseline compared to those who were culture-negative (16% vs 3%; OR=6.9, 95% CI [2.3 – 19.7], p<0.001, see Figure 1). Of all ICU patients who developed *K. pneumoniae* infections and contributed baseline screening swabs, half (n=13/27, 48%) tested positive for *K. pneumoniae* GI carriage at baseline (including eight who were screened >2 days prior to developing the infection).

**Figure 1.**
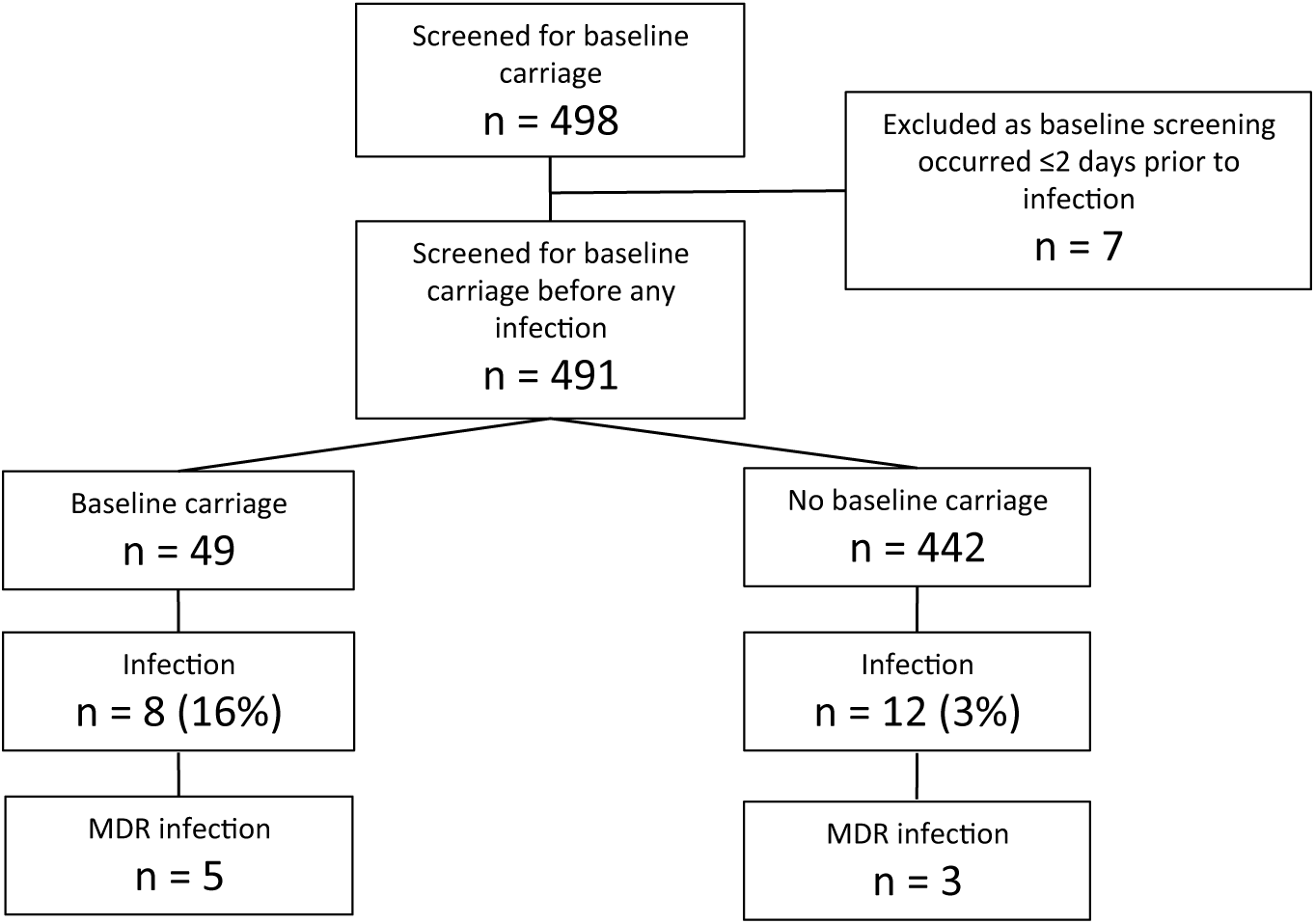
Flowchart outlining number of patients included in each part of rates analyses.

These results show *K. pneumoniae* GI colonisation was significantly associated with *K. pneumoniae* infection in ICU patients. To determine whether infections were caused by patients’ own colonising bacteria and to identify transmission between ICU patients, we sequenced the genomes of all *K. pneumoniae* isolated from patients who had spent any time in the ICU during their hospital stay(s). A total of 143 high quality whole genome sequences were obtained from 106 patients, including 56 clinical, 80 GI carriage and 7 throat carriage isolates. Core genome phylogenetic analysis (Figure 2) revealed the presence of 61 lineages of *K. pneumoniae sensu stricto* (n=111 isolates) and 24 lineages of two closely related species: 20 *K. variicola* lineages (n=28 isolates) and four *K. quasipneumoniae* lineages (n=4 isolates). The latter are recently-described species that are closely related to *K. pneumoniae* and are typically identified as *K. pneumoniae* using biochemical or proteomics-based methods^21^. The average distance between *K. pneumoniae* lineages was 0.5% nucleotide divergence, representing thousands of years of evolutionary separation.

**Figure 2.**
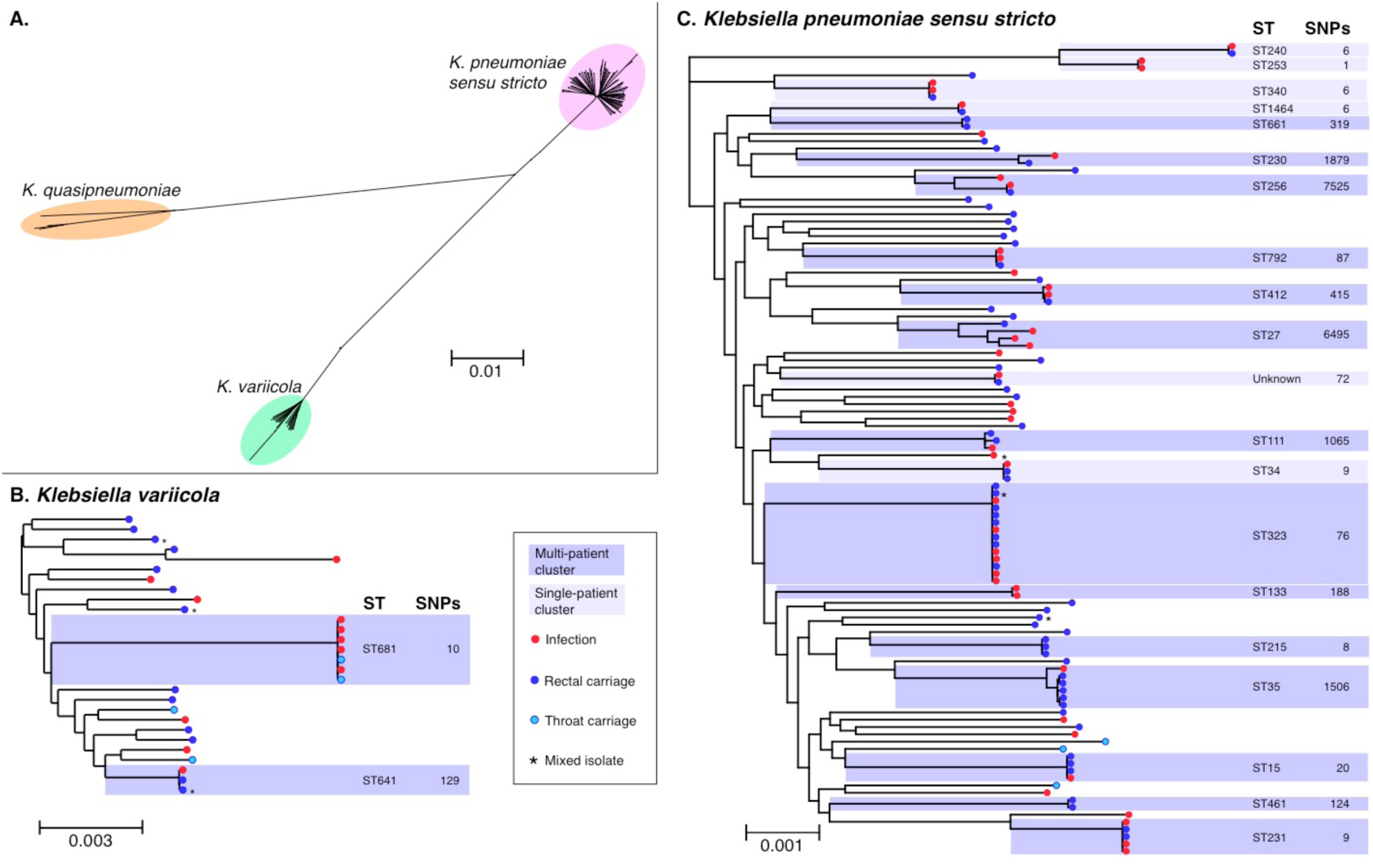
Genome diversity of isolates from ICU patients identified as *Klebsiella pneumoniae*. All trees are maximum likelihood trees inferred from core genome SNP alignments. Scale bars indicate average number of substitutions per site across the genome. Tip colours indicate isolate source as per inset legend. *Mixed isolate, sequence reads include substantial non-*Klebsiella* sequences; position in tree shown is based on *Klebsiella* SNPs. Phylogenetic lineages to which more than one ICU isolate belongs are highlighted and labelled with their corresponding multi-locus sequence type (ST) and the total number of SNPs identified between isolates in the lineage; darker shading indicates multiple patients contributed isolates in that cluster, as per inset legend. (A) Unrooted tree of all isolates, revealing three distinct species that are typically identified as *K. pneumoniae* in diagnostic laboratories. (B) Midpoint rooted species tree for *K. variicola* isolates. (C) Midpoint rooted species tree for *K. pneumoniae sensu stricto* isolates.

Most *Klebsiella* lineages (n=69/85, 81%) were identified in just one patient, and 60% of patients (n=64) had their own unique lineage not observed in any other patients (Figure 2). Half of infections (24/49 patients, 49%) were caused by a lineage unique to that patient. Fifteen patients had both GI carriage and infection isolates available for genome comparison, and 12 of these pairs matched at the lineage level (including six patients whose carriage isolate was collected >2 days prior to the infection). Notably, each of these patients carried their own unique *K. pneumoniae* lineage, except for AH0214 and AH0255 who shared lineage ST323 carriage and infection isolates.

### *Klebsiella* transmission in the ICU

Sixteen *Klebsiella* lineages were detected in more than one patient (dark shading, Figure 2). Lineage sharing between patients could result from recent transmission of bacteria within the hospital (strain sharing), or by independent acquisition of a lineage that has been circulating in the community (lineage sharing). To distinguish these possibilities we compared pairwise SNP distances between isolates from the same patient with those from different patients (Figure 3A). Intra-patient genetic distances were nearly all (97%) less than 25 SNPs per 5 Mbp and most (82%) were less than 10 SNPs, while between-patient genetic distances ranged from 0 to >5,000 SNPs (Figure 3A). Using 25 and 10 SNPs per 5 Mbp as cut-offs to indicate likely and very likely strain sharing between patients, we identified five groups of ICU patients that likely shared *Klebsiella* strains. Strikingly, each of these groups comprised patients with overlapping admissions, making them epidemiologically plausible intra-hospital transmission chains (Figure 4). We identified six patients with infections that were attributable to these intra-hospital transmission chains (n=4 ST681 *(K. variicola)*, n=1 ST323, n=1 ST231; Figure 4), representing 12% (n=6/49) of all ICU patients with infections. These infections included two episodes of pneumonia, two wound infections and two UTIs, one of which disseminated to cause bacteraemia with sepsis. Of the four donors in the transmission chains, three had pneumonia and one had a wound infection. Most of the infections associated with transmission were MDR (3/4 donors and 3/6 recipients), indicating a strong association between MDR infections and transmission in the ICU (OR 13.6, p=0.002). In addition, we observed four patients whose carriage of *K. pneumoniae* was attributable to intra-hospital transmission chains but did not result in any recorded *K. pneumoniae* infection during hospital stay (n=1 ST323 (MDR), n=2 ST215 (non-MDR), n=1 ST15 (MDR)), representing 5% of all patients in whom carriage was detected.

**Figure 3.**
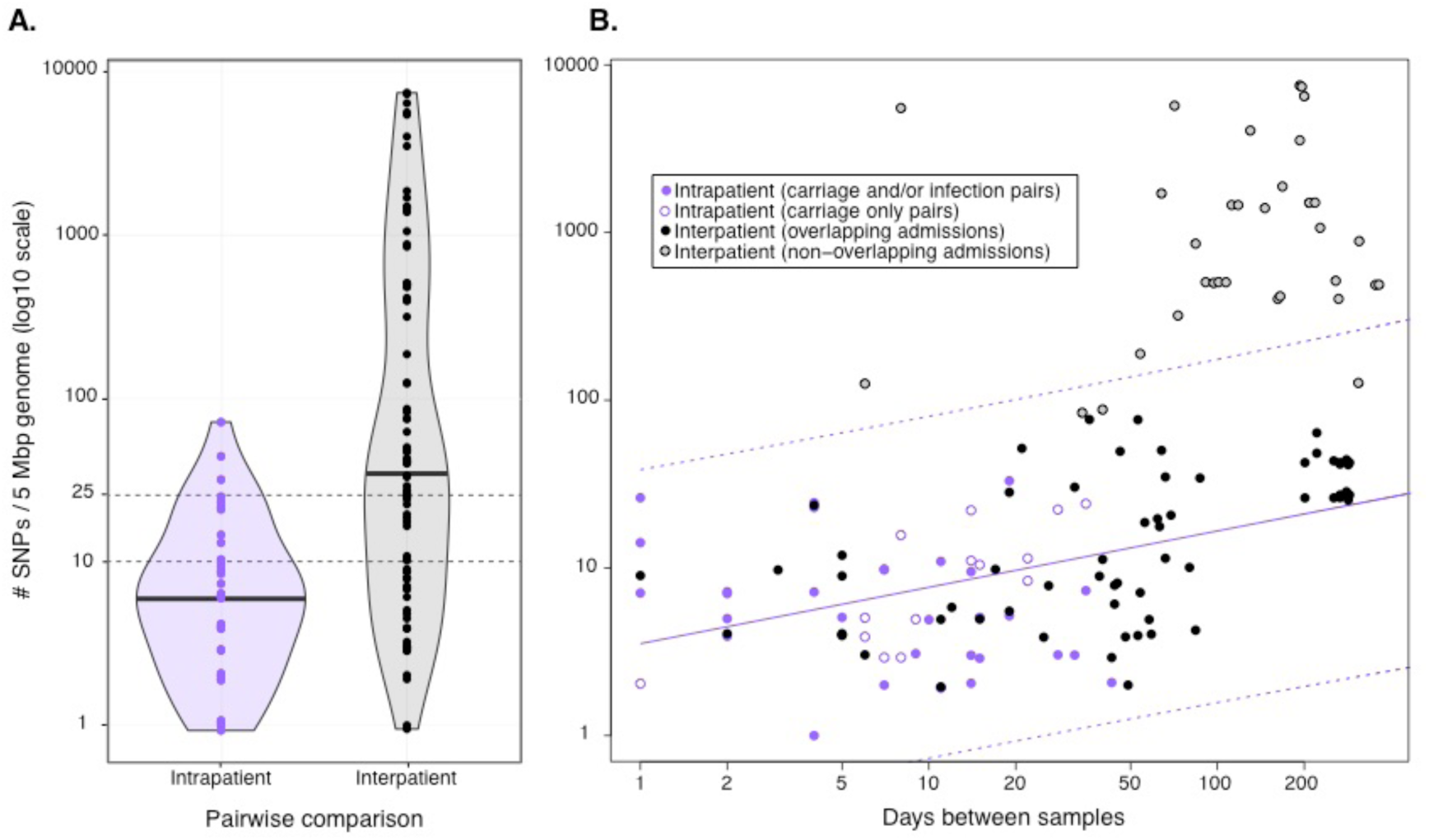
Pairwise genetic distances between isolates belonging to the same lineage, expressed as SNPs per 5 Mbp of genome in order to normalise for differences in shared gene content between strain pairs. (A) Violin plots showing distribution of pairwise genetic distances within and between patients; black bars indicate the median distribution; note log_10_ scale. (B) Scatter plot of pairwise genetic distances (y-axis) on time between isolation (x-axis), coloured according to inset legend; both on log_10_ scale. Solid line, least-squares linear regression line fit to all isolate pairs involved in plausible transmission chains (intra-patient pairs, purple points; and inter-patient pairs associated with overlapping admissions, filled black points); dashed lines indicate 99% prediction interval for the linear model.

**Figure 4.**
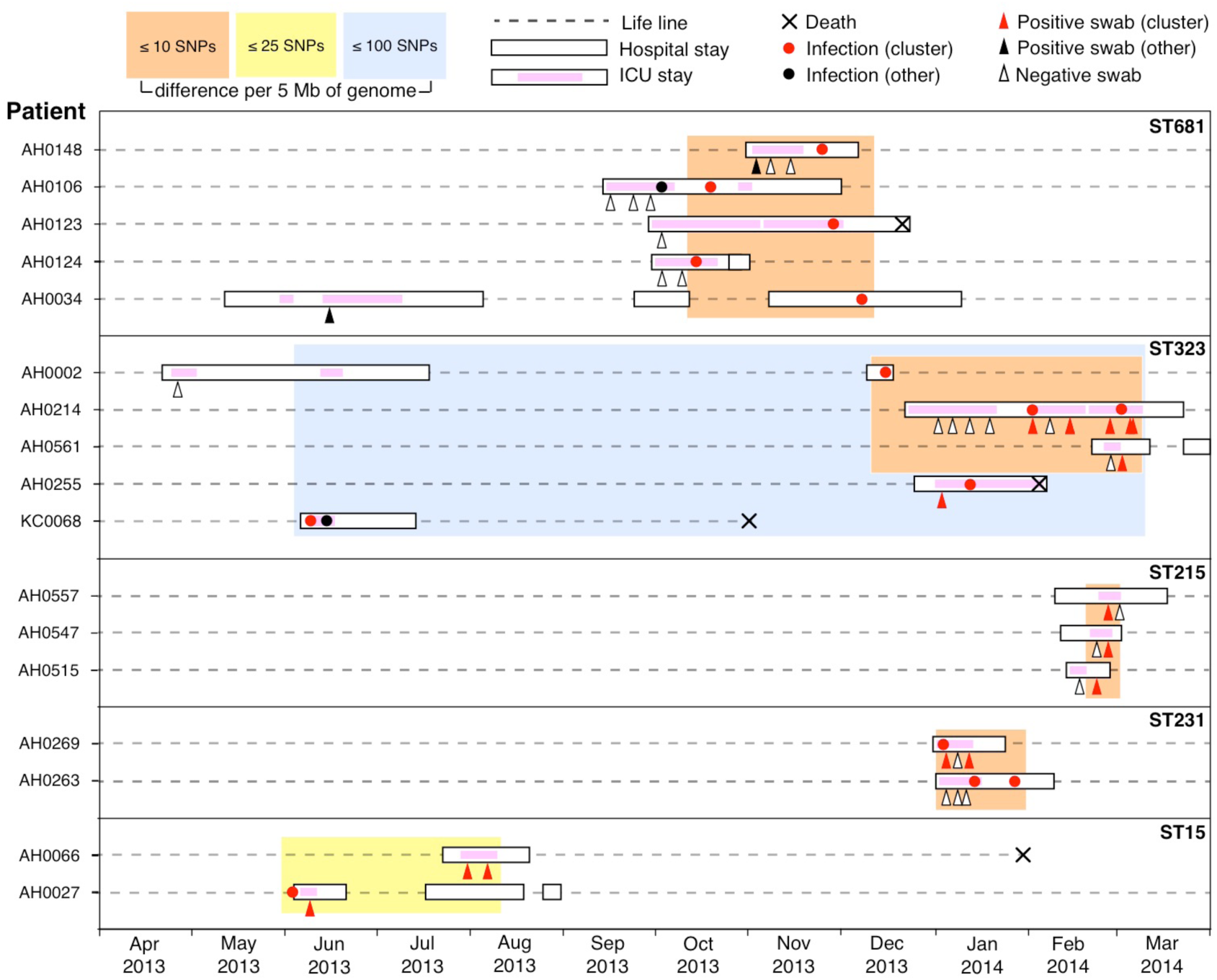
Timelines for all lineages detected in multiple patients that show any inter-patient pairwise genetic distance between isolates of ≤25 SNPs per 5 Mbp. Lineages are boxed and labelled with their multi-locus sequence type (ST). Each horizontal dashed line indicates the time line for a patient, labelled to the left (crosses indicate date of death where applicable). Periods of Alfred Hospital admission are indicated as white boxes, periods in ICU as pink shading. Circles indicate *K. pneumoniae* infection isolates (red, belonging to the lineage; black, other lineage); triangles indicate rectal screening swabs (red, *K. pneumoniae* belonging to the lineage; black, *K. pneumoniae* of another lineage; unfilled, negative for *K. pneumoniae).* Orange boxes indicate groups of isolates for which all patients have at least one pairwise genetic distance of ≤10 SNPs per 5 Mbp with another in the group; similarly for yellow (≤25 SNPs) and blue (≤100 SNPs) boxes.

A further 11 lineages were detected in more than one ICU patient each, but were considered unlikely to represent strain sharing due to excessive pairwise SNP distances. The patients carrying these lineages also lacked overlapping ICU admissions (Supplementary Figure 2). Such lineage sharing could result from independent acquisition in the community prior to hospital admission, but could also potentially be explained by cryptic longer-term transmission chains involving bacteria that persist within the hospital between admissions (for example, in reservoirs such as sinks or drains, or colonisation of healthcare workers^22–24^). In the latter case, we would expect the genetic distances between isolates in these clusters to be consistent with the accumulation of genetic variation over time in an evolving reservoir or transmission chain, which could exceed the 25-SNP cut-off if sufficient time has passed between isolations. To assess this, we compared genetic distances with time-between-isolation for pairs of isolates from the same patients and from within the five epidemiologically supported intra-hospital transmission chains (**Figure 3B**), which showed a significant log-linear relationship (R^2^=0.25, p=4×10^−8^) between genetic distance (SNP accumulation) and time. Observed genetic distances for isolate pairs from the 11 multi-patient lineages with non-overlapping stays exceeded the 99% prediction interval of the linear regression model with the exception of two cases that could potentially reflect plausible cryptic transmission chains (Supplementary Figure 2): (i) patient AH0352’s fourth screening swab was positive for ST641 ten months after the isolation of a ST641 isolate from patient KC0038 at a genetic distance of 125 SNPs per 5 Mbp (value within the 95% prediction interval), and (ii) patient AH0620’s clinical wound isolate was 83 SNPs per 5 Mbp distant from that isolated from patient AH0416 one month earlier (value exceeds the 95% prediction interval, but falls within the 99% prediction interval).

## DISCUSSION

We estimated a 5.9% CA rate for culture-positive GI carriage of *K. pneumoniae*, similar to the 3.9% estimated amongst healthy individuals in the human microbiome project based on 16S rRNA amplicon sequencing of stool samples^10^. HA carriage amongst ICU patients was estimated to be much higher at 19%, with 12% of patients converting from culture-negative at baseline to culture-positive on follow-up. This likely reflects acquisition of bacteria in the hospital (transmission of strains was directly observed in 5% of cases) and/or selection for growth of pre-existing *K. pneumoniae* in the GI microbiome during hospitalisation (MDR rate was 18% in HA baseline carriage and 15% at follow-up, but MDR was not detected in CA baseline carriage). The HA rate estimated here is similar to the culture-positive GI carriage rates estimated in other hospital studies^15,16^. Throat carriage was negligible in our ICU population, much lower than the rates of CA naso/oropharyngeal carriage estimated in both culture-based and culture-free studies^10,12^. This is possibly due to high rates of intubation in ICU patients, which may substantially alter the local microbial communities.

Our data demonstrate that *K. pneumoniae* is a common component of the human GI microbiome and of clinical significance in the ICU setting, as: (i) *K. pneumoniae* carriage on admission to ICU was significantly associated with subsequent *K. pneumoniae* infection (OR 6.9, p=0.0003), consistent with the results reported from the 1970s Denver study (OR 4.0, p=0.0009^15^) and the 2016 Michigan study (OR 4.1, p=0.00002^16^); and (ii) the WGS data confirmed a direct link between colonising and infecting strains in 13 patients (80% of those with paired isolates available for testing), also consistent with the Michigan study^16^. Importantly, we found strong agreement between two methods for estimating the proportion of ICU *K. pneumoniae* infections that are attributable to infection with patients’ own GI microbiota: (i) of all 49 *K. pneumoniae* infections diagnosed in ICU patients during the study period, 49% were associated with *K. pneumoniae* lineages unique to the patient; and (ii) of the 27 *K. pneumoniae* infections diagnosed in ICU patients from whom screening swabs were obtained, 48% occurred in patients who tested positive for prior GI colonisation with *K. pneumoniae.* In contrast, only 12% of infections showed evidence of resulting from intra-hospital transmission in this setting. This suggests that while measures to reduce cross-contamination between patients are necessary, they are not sufficient to eliminate *K. pneumoniae* infections in hospitalised patients, and measures to minimise the risk of infection with the patients’ own microbiome deserves significant attention^25–27^.

Key strengths of this study are the prospective cohort design and the use of WGS to confirm species identification and strain relatedness for all *K. pneumoniae* isolated from ICU patients, regardless of antimicrobial susceptibility. Most previous studies of *K. pneumoniae* colonisation in hospitals have focused on ESBL and/or CP isolates only, and while infections associated with such strains are most complicated to treat, they do not represent the major burden of *K. pneumoniae* infections. WGS demonstrated that some isolates identified as *K. pneumoniae* by MALDI-TOF actually belonged to closely related groups that have recently been described as separate species^21^, *K. quasipneumoniae* and *K. variicola* (Figure 2). Since these species are very closely related to *K. pneumoniae sensu stricto* (˜4% nucleotide divergence), are clinically indistinguishable, and are typically identified as *K. pneumoniae* in diagnostic laboratories, comparable studies reporting *K. pneumoniae* carriage encompass the entire *K. pneumoniae* complex. Hence we included all three species in our summaries of results. WGS also demonstrated that a small number of isolates identified as *K. pneumoniae* were either not *Klebsiella* (n=3), were contaminated with a large proportion of DNA from multiple *Klebsiella* (n=2) or *non-Klebsiella* (n=3), or had very low *Klebsiella* mixture (indicated in Figure 2). These may reflect co-infection followed by selection for different subpopulations in the laboratory used for the different tests, or misidentification by MALDI-TOF. These samples were therefore excluded from high-resolution genomic analysis for attribution purposes (data Figure 1, **Supplementary Table 4**) but were included in calculation of overall rates of infection and carriage, which therefore reflect solely the routine laboratory identification and are directly comparable with other reported results.

The main limitations of the study arise from the swabbing procedure to identify *K. pneumoniae* colonisation on admission. Efforts were made to collect swabs as soon as possible following admission, however the need to obtain informed consent from the participant or a person responsible and not disrupt patient care meant that it was often not possible to obtain screening swabs on the day of admission. Notably, similar culture-positive rates were observed for swabs collected on day 0, 1 or 2 of admission (Supplementary Table 3), so it is unlikely that this had a significant impact on results. We used rectal swab culture to determine *K. pneumoniae* GI colonisation status, however while this approach is standard for pathogen carriage screening^28–30^, its sensitivity to detect *K. pneumoniae* is not well characterised and likely depends on the *K. pneumoniae* strain, GI microbiome composition and recent antimicrobial exposures. There is likely a significant false negative rate, particularly for the detection of CA carriage of susceptible organisms in the absence of recent antibiotic treatment. Hence our study probably underestimates the rate of colonisation in the community and the contribution of colonisation to subsequent infection.

Our conclusion that the GI microbiome is a source of *K. pneumoniae* infections in ICU patients echoes similar findings that colonising strains of *S. aureus, A. baumannii, *Enterococcus*, and Enterobacteriaceae are a common source of HA infections*^16,22–24^. Routine screening for nasal carriage of methicillin resistant *S. aureus* (MRSA) or gut carriage of vancomycin resistant *Enterococcus* (VRE) or ESBL/CP Enterobacteriaceae has been introduced in various hospital settings^28–31^. A recent study of CP *K. pneumoniae* in Israel suggested screening and isolation of carriers could help end current outbreaks and prevent future ones^6^. A similar study introduced screening for EBSL *K. pneumoniae* in order to limit and prevent current and future outbreaks^32^. Whilst those studies focus on screening for CP or ESBL *K. pneumoniae*, our results indicate that routine screening for general *K. pneumoniae* carriage in the ICU could also be a valuable tool. In particular, foreknowledge of the antimicrobial susceptibility profiles of *K. pneumoniae*, as well as other opportunistic pathogens resident in the microbiome, could guide the choice of prophylactic and therapeutic antimicrobial treatment^25,26,28^ .

## Acknowledgements

This work was funded by the NHMRC of Australia (project #1043822 and Fellowship #1061409 to K. E. H) and C. L. G. was supported by an Australian Government Research Training Program Scholarship. The authors gratefully acknowledge the contribution and support of Janine Roney, Mellissa Bryant, Jennifer Williams, Iain Abbott and Noelene Browne at the Alfred Hospital, and the sequencing team at the Wellcome Trust Sanger Institute.

---

**Supplementary Table 1. Carriage and infection sample data and accession numbers.** [see supplementary document called SupplementaryTable1_SampleDataAndAccessionNumbers.csv]

**Supplementary Table 2.** *K. pneumoniae* carriage rates detected from baseline swabs of ICU patients. Last two columns show comparison of rates in females vs males: OR, odds ratio; P, p-value.

| Age | Total | Carriage<br>N (%) | Female<br>carriers<br>N (%) | Male<br>carriers<br>N (%) | Association with female sex<br>(Fisher's exact test) |  |
| --- | --- | --- | --- | --- | --- | --- |
|  |  |  |  |  | OR (95% CI) | p-value |
| ≤25 | 28 | 3 (11%) | 1 (14%) | 2 (10%) | 1.6 (0.0, 35.2) | 1 |
| 26-35 | 44 | 5 (11%) | 1 (6%) | 4 (15%) | 0.3 (0.0, 3.8) | 0.6 |
| 36-45 | 47 | 4 (9%) | 1 (5%) | 3 (8%) | 0.4 (0.0, 4.8) | 0.6 |
| 46-55 | 85 | 5 (6%) | 1 (4%) | 4 (7%) | 0.6 (0.0, 6.7) | 1 |
| 56-65 | 107 | 9 (8%) | 3 (10%) | 6 (8%) | 1.4 (0.2, 7.0) | 0.7 |
| 66-75 | 89 | 11 (12%) | 3 (10%) | 8 (14%) | 0.7 (0.1, 3.3) | 0.7 |
| 76-85 | 77 | 15 (19%) | 7 (25%) | 8 (14%) | 1.7 (0.5, 6.2) | 0.4 |
| >85 | 21 | 2 (10%) | 0 (0%) | 2 (13%) | 0 (0.0, 13.8) | 1 |
| **All** | **498** | **54 (10.8%)** | **17 (10%)** | **37 (11%)** | **0.9 (0.5, 1.8)** | **0.9** |

**Supplementary Table 3.** Baseline GI carriage rates, with CA cohort broken down into individual days.

|  | Total swabbed (n) | Kp positive (%) | Kp negative (%) |
| --- | --- | --- | --- |
| **CA / Day 0-2** | **324** | **19 (5.9%)** | **305 (94.1%)** |
| - Day 0 | 24 | 1 (4%) | 23 (96%) |
| - Day 1 | 206 | 15 (7%) | 191 (93%) |
| - Day 2 | 94 | 3 (3%) | 91 (97%) |
| **HA / Day 3+** | **174** | **33 (19.0%)** | **141 (81.0%)** |
| **Total** | **498** | **52 (10.4%)** | **446 (89.6%)** |

**Supplementary Table 4. Patients with time in the ICU, with carriage isolates, infection isolates, or both carriage and infection isolates.**

[see supplementary document called SupplementaryTable4_ICUpatientData.csv]

**Supplementary Figure 1.**
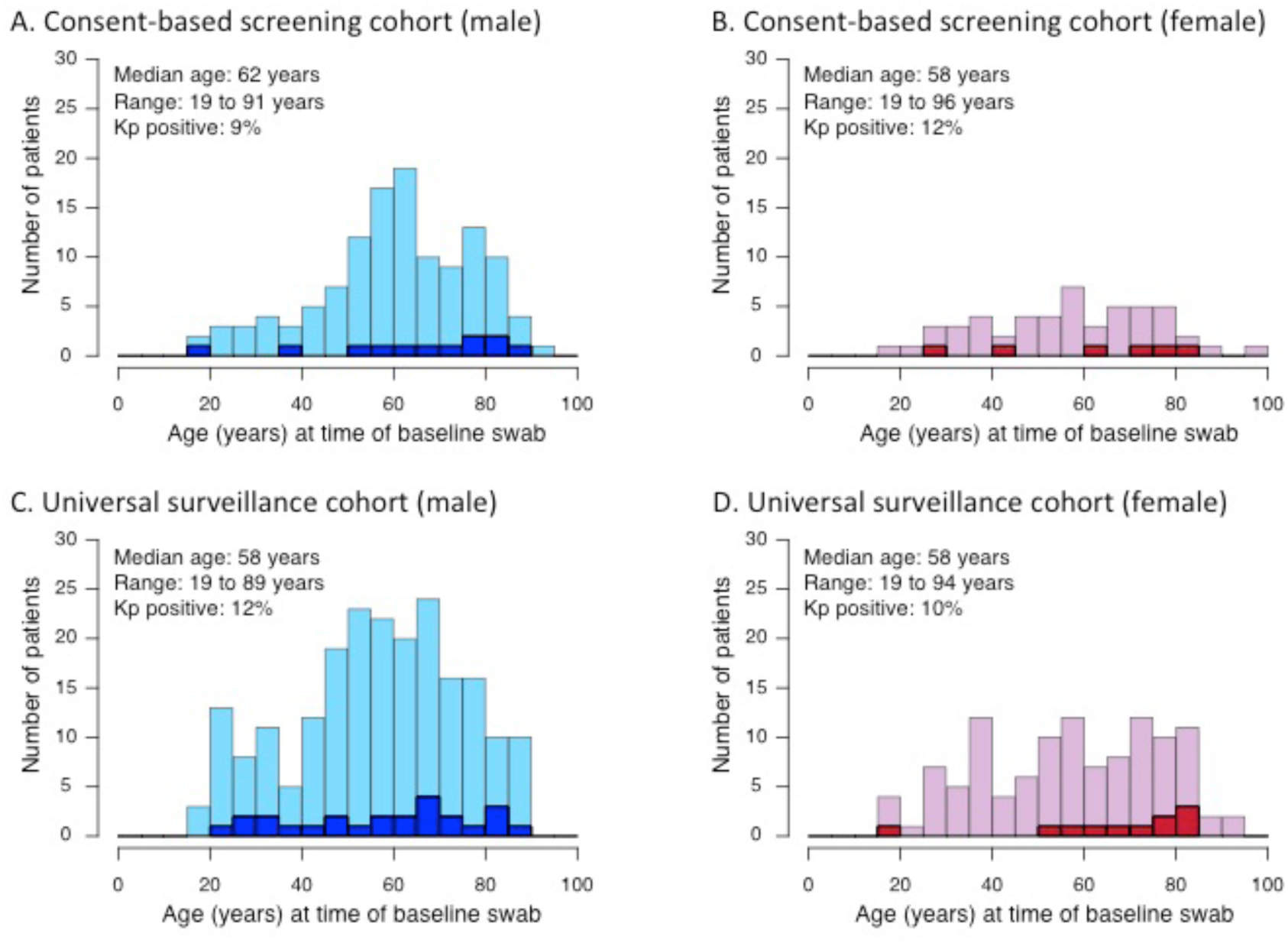
Distribution of age and *K. pneumoniae* culture positive rates, by gender, among individuals screened for baseline carriage of *K. pneumoniae* over the first nine months (panels A and B) and the final three months (panels C and D) of the study period. Lighter colours, culture negative individuals; darker colours, culture positive.

**Supplementary Figure 2.**
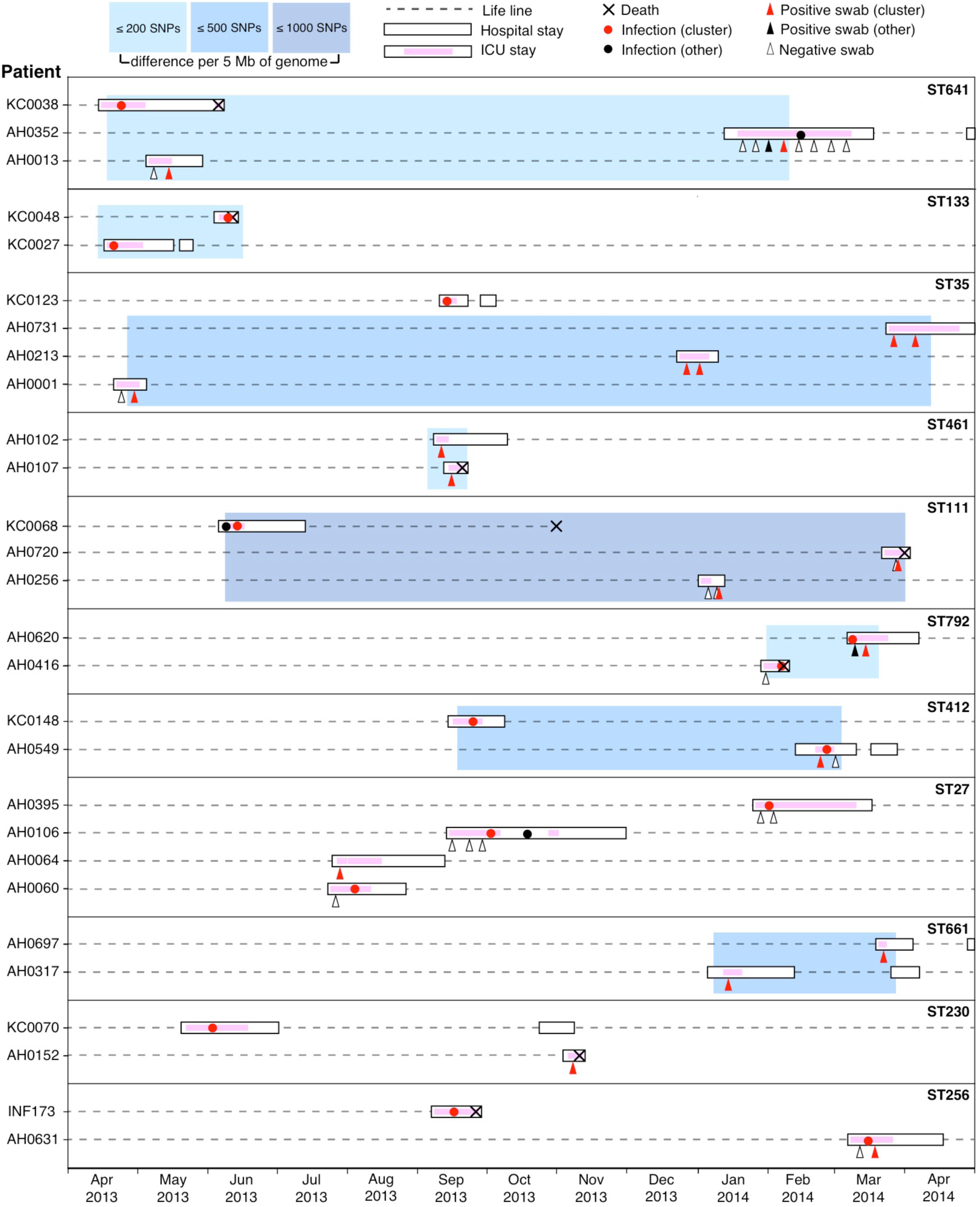
Timelines for all lineages detected in multiple patients that did not show any inter-patient pairwise genetic distance between isolates of ≤25 SNPs per 5 Mbp. Lineages are boxed and labelled with their multi-locus sequence type (ST). Each horizontal dashed line indicates the time line for a patient, labelled to the left (crosses indicate date of death where applicable). Periods of Alfred Hospital admission are indicated as white boxes, periods in ICU as pink shading. Circles indicate *K. pneumoniae* infection isolates (red, belonging to the lineage; black, other lineage); triangles indicate rectal screening swabs (red, *K. pneumoniae* belonging to the lineage; black, *K. pneumoniae* of another lineage; unfilled, negative for *K. pneumoniae).* Blue boxes indicate groups of isolates for which all patients have at least one pairwise genetic distance from another in the group that falls below the cut-off indicated in the inset legend.

